# Global neural encoding of model-free and inference-based strategies in mice

**DOI:** 10.1101/2024.02.08.579559

**Authors:** Shuo Wang, Huayi Gao, Kotaro Ishizu, Akihiro Funamizu

**Affiliations:** Institute for Quantitative Biosciences, University of Tokyo, Laboratory of Neural Computation, 1-1-1 Yayoi, Bunkyo-ku, Tokyo 113-0032, Japan; Department of Life Sciences, Graduate School of Arts and Sciences, University of Tokyo, 3-8-2, Komaba, Meguro-ku, Tokyo 153-8902, Japan

## Abstract

When a simple model-free strategy does not provide sufficient outcomes, an inference-based strategy estimating a hidden task structure becomes essential for optimizing choices. However, the neural circuitry involved in inference-based strategies is still unclear. We developed a tone frequency discrimination task in head-fixed mice in which the tone category of the current trial depended on the category of the previous trial. When the tone category was repeated every trial, the mice continued to use the default model-free strategy, as well as when tone was randomly presented, to bias the choices. In contrast, the default strategy gradually shifted to an inference-based strategy when the tone category was alternated in each trial. Brain-wide electrophysiological recording during the overtrained phase suggested that the neural activity of the frontal and sensory cortices, hippocampus, and striatum was correlated with the reward expectation of both the model-free and inference-based strategies. These results suggest the global encoding of multiple strategies in the brain.

## Introduction

Perceptual decision-making requires estimating a hidden context from the observation of sensory inputs. Signal detection theory (SDT) shows that to optimize behavior, subjects also need to infer expected outcomes in each context (value) and how the context changes over time (context transition probability)^1,2^. This study investigated how various brain regions represent decision-making based on value or context probability.

One simple strategy for optimizing behavior by estimating the expected outcome (value) is to estimate and update the value of each choice through trial and error from past direct experiences. This is accomplished by model-free reinforcement learning (RL)^3–5^. Model-free RL does not estimate the transition of context when making choices^3^. However, as noted in previous theoretical^6^ and experimental research^7–9^, model-free RL is not always the best strategy for optimizing choices. For example, when contexts have certain dependencies or structures, a simple RL model involving only value estimation fails to optimize choices^10–12^. In such complex environments, a behavioral strategy in which the hidden structure of context relationships is inferred becomes important^13^. This is also supported by the SDT, as both value and context estimations are essential for optimizing behavior^2^. A strategy using an internal model of context-transition probability is named the abstract state-based model^7^ or inference-based strategy^8^.

Previous experiments in rodents, monkeys, and humans have shown that the cortico-basal ganglia circuit, which includes the striatum (STR), motor cortex, prefrontal cortex, and sensory cortex, is involved in model-free RL^4,14–20^. In contrast with the model-free strategy, the neural circuit of the inference-based strategy is still under investigation. Early human studies utilizing sophisticated behavioral tasks have identified parallel pathways for model-free and inference- or model-based strategies in the brain^21,22^, while others have shown overlapping involvement of brain regions in these two strategies^23^. These studies revealed that the prefrontal cortex is involved in inference-based strategies^21–23^. Recently, rodent studies have shown that the orbitofrontal cortex (OFC) and hippocampus (HPC) are necessary for the inference strategy^8,24^. In the early phase of training, mice use model-free RL as a default strategy. Mice then change to an inference-based strategy at the late phase of training when the simple default strategy does not provide sufficient outcomes in a foraging task^8^. Although the neural circuits involved in the inference-based strategy are gradually being identified in some brain regions in animal experiments, it is unclear how various regions in the brain represent the inference-based strategy and whether the circuits are distinct or similar to those involved in the model-free strategy.

Here, we updated our previous tone frequency discrimination task^1,25–27^ to test the neural representations of model-free and inference-based strategies. The task probabilistically alternated the tone category of the current trial based on the category in the previous trial with a transition probability of *p*^28,29^. We first trained all the mice in the neutral condition (*p* = 0.5), where there was no bias of tone presentation in the task. We found that although the optimal behavior was an unbiased selection of the left or right choice, mouse behavior was already biased by the outcome in previous trials, suggesting that the default strategy of mice was value-based model-free RL. We then divided the mice into two groups: one group repeated one tone category (repeating condition: *p* = 0.2), while the other group alternated the tone category in every trial (alternating condition: *p* = 0.9). Interestingly, the acquisition of proper choice biases was faster in the repeating condition than in the alternating condition. Biased behavior in the repeating condition was achieved from the first session, suggesting that the default model-free strategy was used to optimize choices. In contrast, the acquisition of choice biases in the alternating condition took 3 sessions to achieve, and the behavior was gradually fit to a state-based model. Value updating was observed even at the overtrained phase in the repeating condition, while the choice was stable in the alternating condition, further supporting that different strategies were utilized in the two conditions.

We obtained brain-wide electrophysiological recordings from the OFC, HPC, STR, primary motor cortex (M1), posterior parietal cortex (PPC), and auditory cortex (AC) during the overtrained phase of the task. We found that, in both conditions, the neurons in all the recorded regions showed increased activity when the choice was expected to have a high reward probability^30^. In contrast, at the outcome timing, the neurons increased the activity with unexpected outcomes mainly in the repeating condition, possibly because the behavior in the alternating condition was already stable during electrophysiology and did not need to update the choices based on outcomes. These results suggest the global encoding of multiple strategies in the brain.

## Results

### Mouse choices depend on the transition of tone category in a tone frequency discrimination task

In our tone frequency discrimination task, mice were head-fixed and placed on a treadmill^1,25,31^ (**Fig. 1a, top**). Each trial began with retracting the spouts away from the mouse. After a random interval of 1.0–2.0 s, a tone stimulus with a duration of 0.6 s was presented from a speaker placed to the right front of the mouse (**Fig. 1a, bottom left**). The tone stimuli were tone clouds, which were mixtures of low-frequency (5–10 kHz) and high-frequency (20–40 kHz) pure tones (**Fig. 1a, bottom right**)^1,25–27^. Depending on the dominant frequency, tone clouds were categorized as low or high. In addition, the tone clouds were named easy (0% and 100% of high-frequency tones), moderate (20%/80% or 25%/75% of high-frequency tones), or difficult (35%/65% or 45%/55% of high-frequency tones). The association between the tone category (low or high) and correct choice was determined for each mouse. A correct or incorrect choice resulted in the provision of 10% sucrose water (2.4 µl) or a noise burst (0.2 s), respectively.

**Fig. 1.**
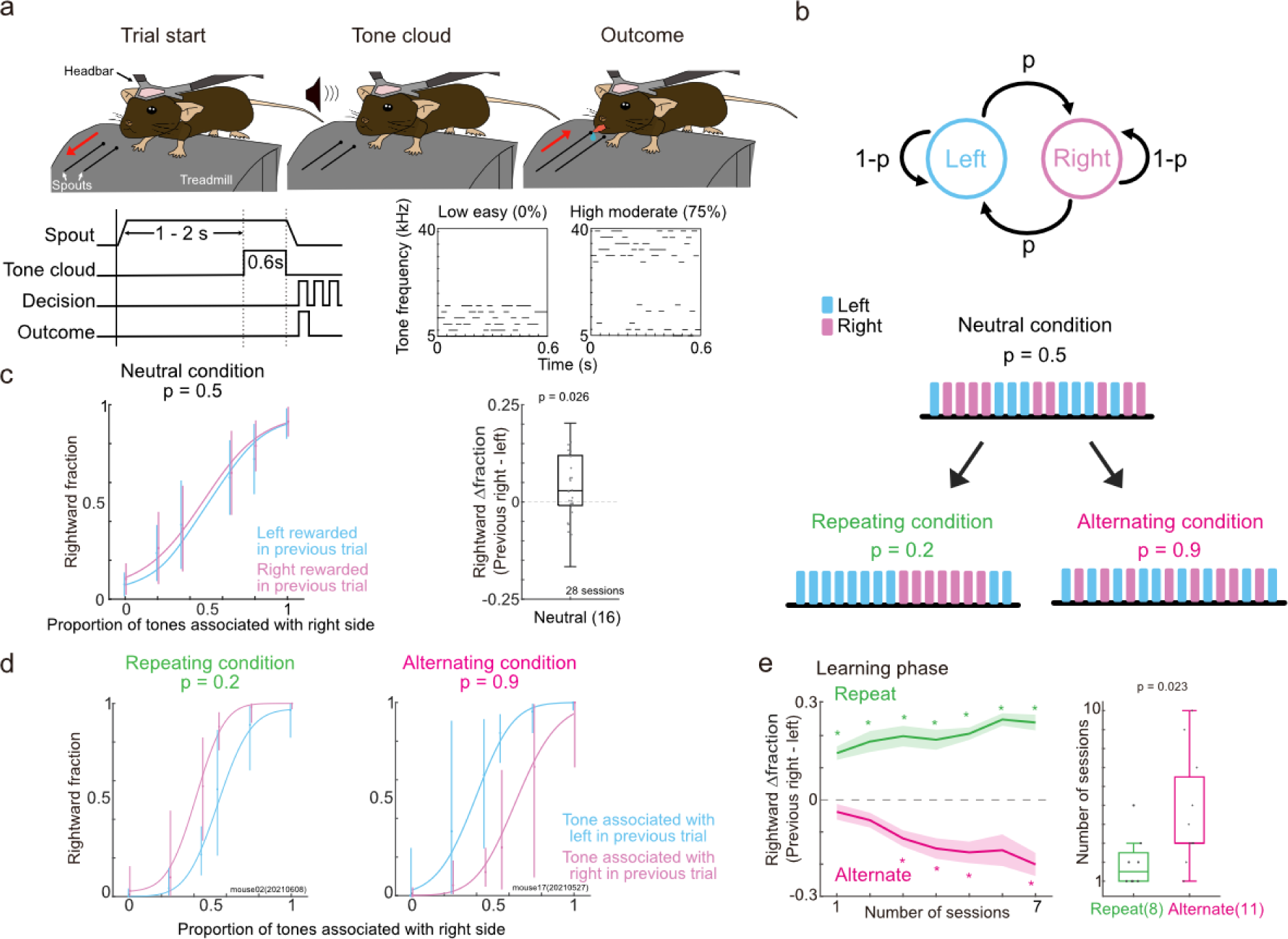
Tone frequency discrimination task in head-fixed mice with repeating and alternating conditions. **a.** Task scheme. Each trial started by moving the spouts away from the mouse. After a random interval of 1–2 s, a sound stimulus (tone cloud) was presented from the speaker positioned in the right front of he mouse. The spouts immediately approached the mice after the end of the sound. Mice licked either he left or right spout to receive sucrose water. The right panels show example tone clouds. During the overtrained electrophysiological phase, the spouts were moved 0.5 s after the end of the sound. **b.** Task conditions. The task included neutral, repeating, and alternating conditions. After the neutral condition with a transition probability of *p* = 0.5, mice were exposed to either the repeating condition (*p* = 0.2) or the alternating condition (*p* = 0.9). **c.** Means and standard deviations of psychometric function in the neutral condition (*p* = 0.5) (28 sessions in 16 mice). Mice significantly biased their choices to the previously rewarded side (linear mixed-effects model) (central mark in the box: median, edges of the box: first quartile (Q1) and third quartile (Q3); bars: most extreme data points without outliers, here and hereafter). **d.** Example psychometric function of choice behavior in the repeating (left) and alternating conditions (right). Error bars are the 95% confidence intervals. **e.** Comparison of the number of sessions required to achieve the proper choice biases in the repeating and alternating conditions after switching from the neutral condition. (Left) Rightward Δ fraction was the difference in the average raction of right-side choice after the right- and left-rewarded trials. Means and standard errors repeating condition: 56 sessions in 8 mice; alternating condition: 77 sessions in 11 mice, * p < 0.01 in he Wilcoxon signed rank test). (Right) The number of sessions required to achieve the proper choice biases (**Methods**) (8 and 11 mice; Mann‒Whitney U test).

In our task, the tone category of the current trial was probabilistically alternated based on the tone category in the previous trial with a transition probability of *p* (**Fig. 1b**)^28,29^. We first exposed all the mice to the neutral condition (*p* = 0.5), in which the tone categories were randomly presented. We then divided the mice into two groups. In the repeating condition (*p* = 0.2), tone categories frequently repeated across trials, while in the alternating condition (*p* = 0.9), tone categories alternated across trials.

In the neutral condition, although the tone category was randomly selected in each trial, the choice behavior of the mice was biased toward the side that was rewarded in the previous trial (**Fig. 1c**), suggesting that the mice had a default strategy to repeat the previously rewarded choice. We then analyzed the choices of mice in the repeating and alternating conditions (repeating condition, 113 sessions in 8 mice; alternating condition, 141 sessions in 11 mice). In the example repeating condition, the mouse chose the right side more frequently after trials in which right-side rewarded tones were used than after trials in which the left-side rewarded tones were used, indicating repeating choice biases (**Fig. 1d**). On the other hand, in the alternating condition, the mouse tended to switch choices. To quantify choice bias, we analyzed how the choice in the current trial depended on the tone category in the previous trial. When the task condition switched from the neutral to the repeating condition, the mice immediately showed repeating choice bias from the first session. In contrast, mice in the alternating condition required 3 sessions on average to acquire the alternating biased behavior (**Fig. 1e, Supplementary Fig. 1c and d**). Since mice had repeating choice biases in the neutral condition, these results suggest that mice continuously used the same default strategy to optimize choices in the repeating condition, while mice required some sessions to switch the choice biases in the alternating condition.

### Mice used model-free and inference-based strategies in the repeating and alternating conditions, respectively

We investigated the behavioral strategies of mice under the repeating and alternating conditions. We proposed two models to analyze mouse choices based on the SDT, which predicted that estimations of expected outcomes and hidden-context probabilities are essential for optimizing behavior (**Fig. 2a**)^1,2^. Model-free RL estimated the expected choice outcome in the low- and high-tone-category states, while the belief probabilities of right- and left-rewarded states were constant. The state-based model estimated the probability of current context as the prior belief state. The prior belief state was estimated from the previous state and the state-transition probability, which was updated by a state prediction error in each trial (**Methods)**^32^. In example sessions, the simulated choices generated by the model-free RL and state-based models captured the repeating and alternating choice biases in the repeating and alternating conditions, respectively (**Fig. 2b and c**).

**Fig. 2.**
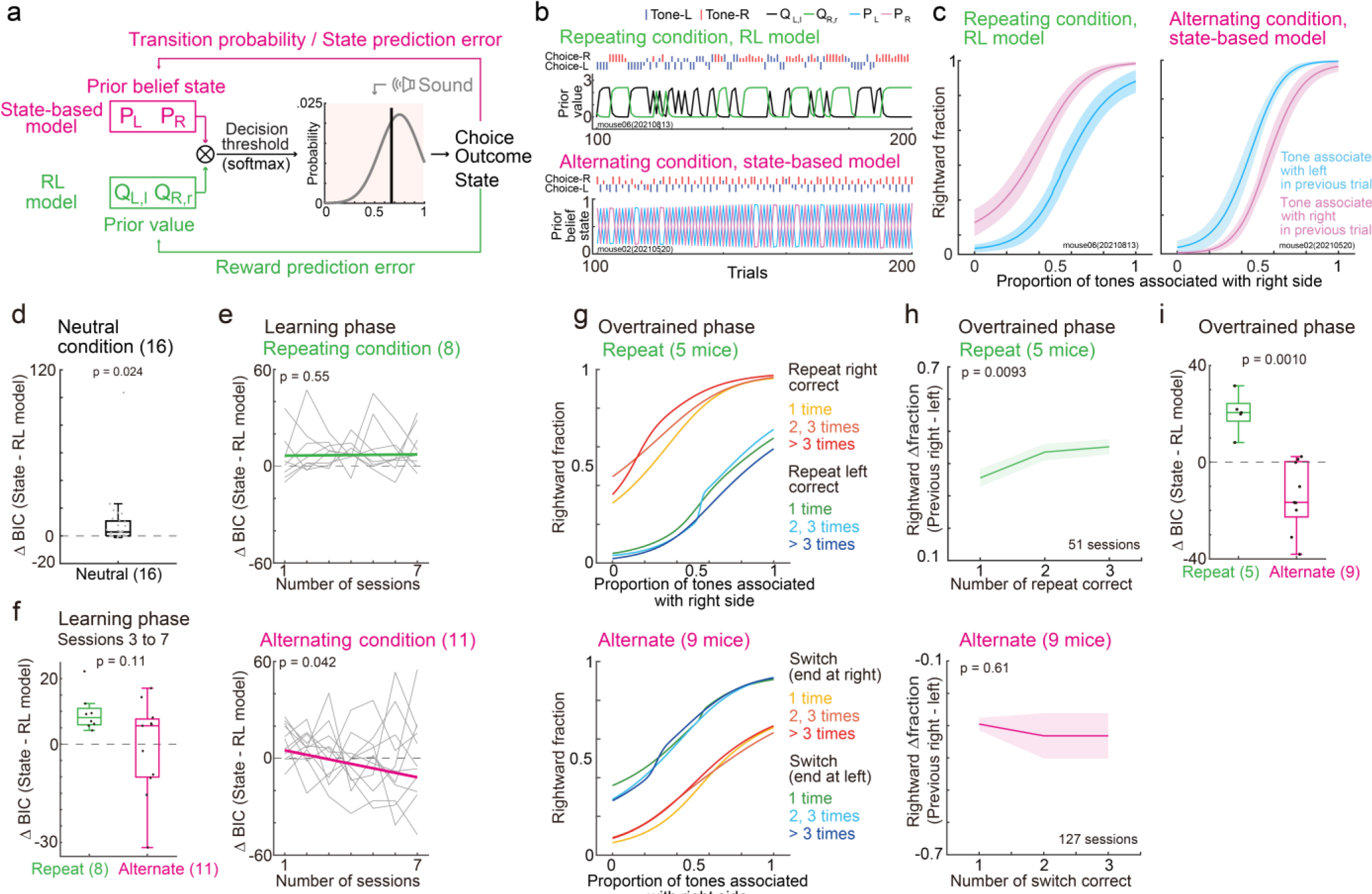
Mice used model-free and inference-based strategies in the repeating and alternating conditions. **a.** Scheme of the reinforcement learning (RL) and state-based models. **b**. Left and right prior values (QL,l and QR,r) were estimated from the RL model in an example session of the repeating condition (top). Prior belief states (PL and PR) were estimated from the state-based model in the alternating condition (bottom). **c.** Example sessions with simulated choices with the RL model in the repeating condition (left) and with the state-based model in the alternating condition (right). We simulated the mice choices 100 times based on the fitted parameters in the RL and state-based models. Means and standard deviations. **d.** Model fitting in the neutral condition. Δ BIC (Bayesian information criterion) was the difference of fitting between the state-based and RL models (28 sessions in 16 mice, linear mixed-effects model). **e.** Model fitting in sessions 1 to 7 in the repeating and alternating conditions (133 sessions in 19 mice, Wilcoxon signed rank test). **f.** Comparison of Δ BIC between the repeating and alternating conditions from sessions 3 to 7 during the learning phase (8 and 11 mice, p value in the Mann‒Whitney U test). **g.** Average psychometric functions of choice behavior in the repeating (51 sessions in 5 mice) and alternating conditions (127 sessions in 9 mice) at the overtrained phase. The transition probability of tone category in the alternating condition was set to *p* = 0.8. **h.** Rightward Δ fractions increased when the number of repeated correct choices increased in the repeating condition. Rightward Δ fractions remained stable in the alternating condition (178 sessions in 14 mice, p value in the linear-mixed effects model). Means and standard errors. **i.** Comparison of Δ BIC between the repeating and alternating conditions during the overtrained phase (5 and 9 mice, p value in the Mann‒Whitney U test).

We first analyzed the choices in the neural condition. We found that the RL model matched the mouse choices better than the state-based model did, suggesting that the mice used a model-free strategy as the default strategy (**Fig. 2d**). In the repeating condition, the RL model was a more consistent fit for choice behavior than the state-based model beginning from the first session. In contrast, in the alternating condition, the choices of the mice gradually fit the state-based model **(Fig. 2e**). When we analyzed only the sessions exceeding 80% accuracy for easy tones, the state-based model matched the mouse choices better than the RL model did in the alternating condition, while the RL model matched the choices in the repeating condition (**Supplementary Fig. 1f**). These results suggest that while the default model-free strategy was used to optimize choices in the repeating condition, mice gradually shifted to an inference-based strategy in the alternating condition.

We also analyzed behavior in the overtrained phase, in which we electrophysiologically recorded the neural activity of the mice. We analyzed 178 sessions in 14 mice during the overtrained phase. In the repeating condition (51 sessions in 5 mice), when the number of correct repeated choices increased, the choice biases increased (**Fig. 2h, top**). This finding suggested that the reward expectation was updated even during the overtrained phase. In contrast, in the alternating condition (127 sessions in 9 mice), the alternating choice biases did not depend on the number of alternated correct choices (**Fig. 2h, bottom**). The RL and state-based models matched the mice’s choices in the repeating and alternating conditions, respectively (**Fig. 2i**), further supporting that mice used model-free and inference-based strategies in the repeating and alternating conditions, respectively.

### Choice and reward are widely encoded in the brain, and only the AC encodes sound

To compare the neural encoding of the model-free and inference-based strategies in the repeating and alternating conditions, we used a Neuropixels 1.0 probe to electrophysiologically record neural activity at the overtrained phase of task (**Fig. 3a**). The transition probability of tone category was set to *p* = 0.2 and *p* = 0.8 in the repeating and alternating conditions, respectively. We targeted the OFC, PPC, HPC, and AC in both the repeating and alternating conditions and additionally recorded the activity of M1 and STR in the alternating condition (**Fig. 3a and b, Supplementary Figs. 2 and 3a**). We used one Neuropixels probe in each session and analyzed the activity of 14 mice in 178 sessions. After spike sorting, we identified 27668 neurons in total (**Methods**).

**Fig. 3.**
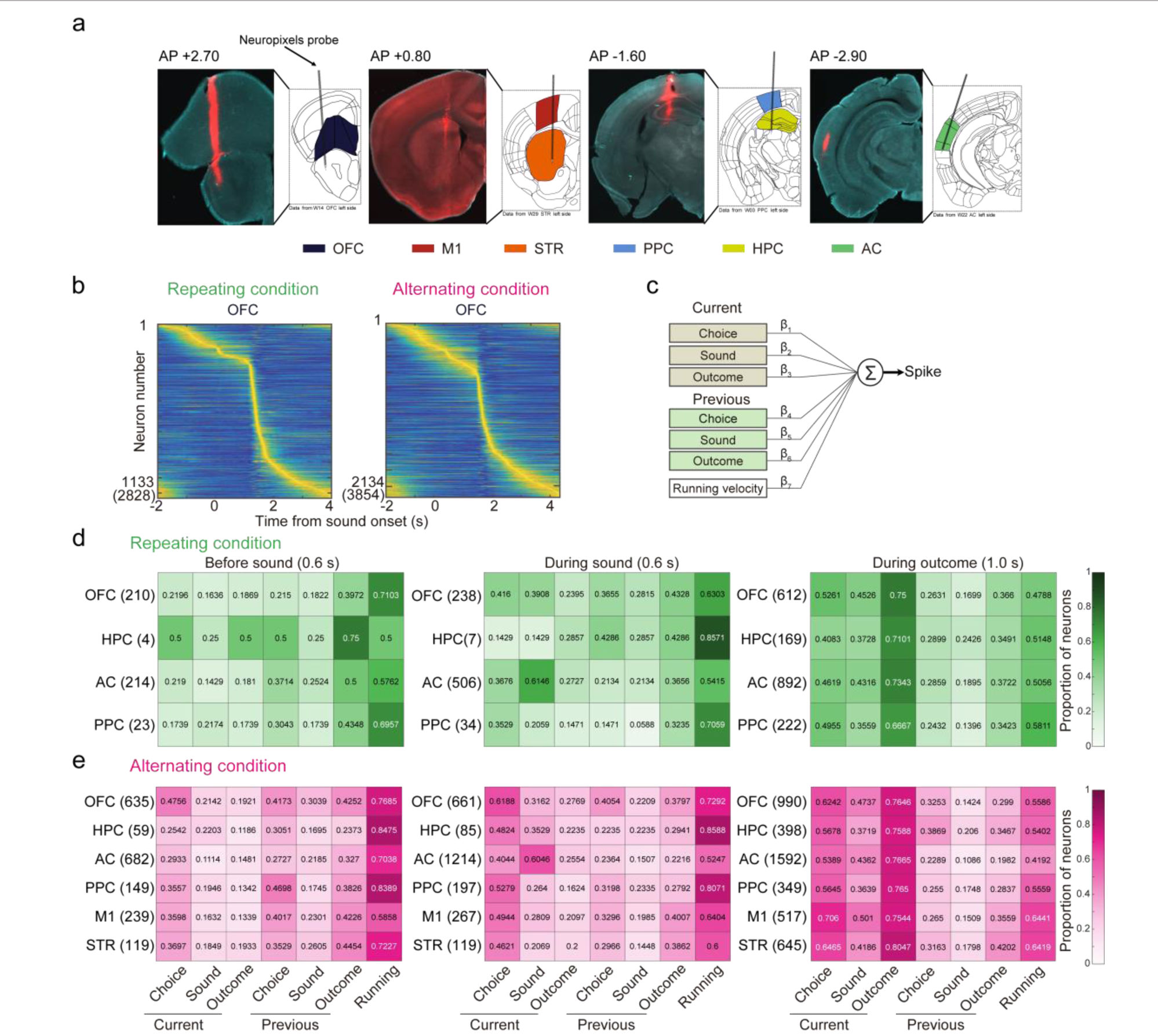
Choice, sound, and outcome representations in the OFC, M1, STR, PPC, HPC, and AC. **a.** Neuropixels 1.0 probe traces for the OFC, M1, STR, PPC, HPC, and AC in example mice. The red color shows the location of the Neuropixels 1.0 probe. **b.** Average activity of task-relevant neurons in the OFC in the repeating (left) and alternating conditions (right) (p < 1.0e-10 in the one-sided Wilcoxon signed rank test). Activity of each task-relevant neuron was normalized between 0 and 1, sorted by the maximum activity timing. The parentheses show the number of all the recorded neurons. **c.** Schematic of regression analysis for testing neural encoding. **d, e.** Proportion of neurons representing the sounds, choices, and outcomes at previous and current trials. Numbers in the parentheses show the number of the subset of task-relevant neurons that showed a significant increase in activity during each time window.

We first detected 12749 task-relevant neurons that exhibited significantly increased activity compared to the baseline activity in one of the 70 time windows during the task (p < 1.0e-10 in the one-sided Wilcoxon signed rank test; 46.08% of all recorded neurons; repeating condition, 3599 out of 9514 neurons (37.83%); alternating condition, 9150 out of 18154 neurons (50.40%)) (**Fig. 3b**, **Supplementary Fig. 3a and b**). The duration of each window was 0.1 s between −1.5 and 2.5 s from sound onset (40 windows). The time windows were also set between −0.5 and 2.5 s from the choice timing (30 windows; 40 + 30 = 70 windows in total). The baseline for sound-onset activity was defined as the activity at −0.2–0 s from the start of the trial, i.e., spout removal. The baseline for choice activity was defined as the activity at −0.2–0 s from the time the spout approached, i.e., between the end of sound and the choice. Among the task-relevant neurons, we targeted the neurons that showed increased activity (1) before sound (−0.6–0 s from the sound onset), (2) during the sound (0–0.6 s from the sound onset), and (3) during the choice and outcome (0–1.0 s from the choice) (**Methods**).

Regression analysis was used to investigate the neural encoding of task variables, including choices, sounds, and outcomes, in the current or previous trials in addition to running speed (**Fig. 3c**) (**Methods**). Before sound onset, the proportions of neurons in each brain region representing the choice in the previous trial were 21.50–50.00% and 27.27–46.98% in the repeating and alternating conditions, respectively **(Fig. 3d and e)**. More neurons represented the choice in the previous trial than the previous sound in multiple brain regions (chi-square test: OFC, PPC, M1, p = 2.6e−04–8.5e−04; HPC, AC, STR, p = 0.032–0.20). During the sound, the proportion of sound-representing neurons in the AC was greater than that in the other regions (AC: 60.46 and 61.46% in the repeating and alternating conditions; OFC, PPC, HPC, M1, STR: 14.29–39.08%; p = 3.5e-20–0.0010 in the chi-square test). At the time of choice, all 6 brain regions were more likely to represent the outcome than the choice or sound (chi square test; outcome vs. choice: AC, OFC, PPC, HPC, STR, p = 1.3e-26–8.6e−04; M1, p = 0.36; outcome vs. sound: all 6 regions: p = 2.7e-47–2.7e−07). These results suggest that the neurons in the AC represent choices, sounds, and outcomes, whereas the neurons in the other brain regions represent choices and outcomes.

### Neurons in wide brain regions modulate activity according to previous and upcoming choices before sound onset

We first analyzed neural activity before sound onset (**Fig. 4a**). We identified 2259 neurons that exhibited a significant increase in activity between −0.6 and 0 s from sound onset (451 and 1883 neurons in the repeating and alternating conditions, respectively; 12.53% and 20.58% of the task-relevant neurons). In the repeating condition, the example neuron in the OFC showed increased activity when the previous trial was a left-rewarded choice followed by a left choice in the current trial (**Fig. 4b, top**). In the alternating condition, an OFC neuron showed increasing activity in the previous right-rewarded trials and the current left-choice trials (**Fig. 4b, bottom**). Thus, example neurons in the OFC showed increased activity in response to the correct choice in previous trials before sound onset (**Fig. 4b**).

**Fig. 4.**
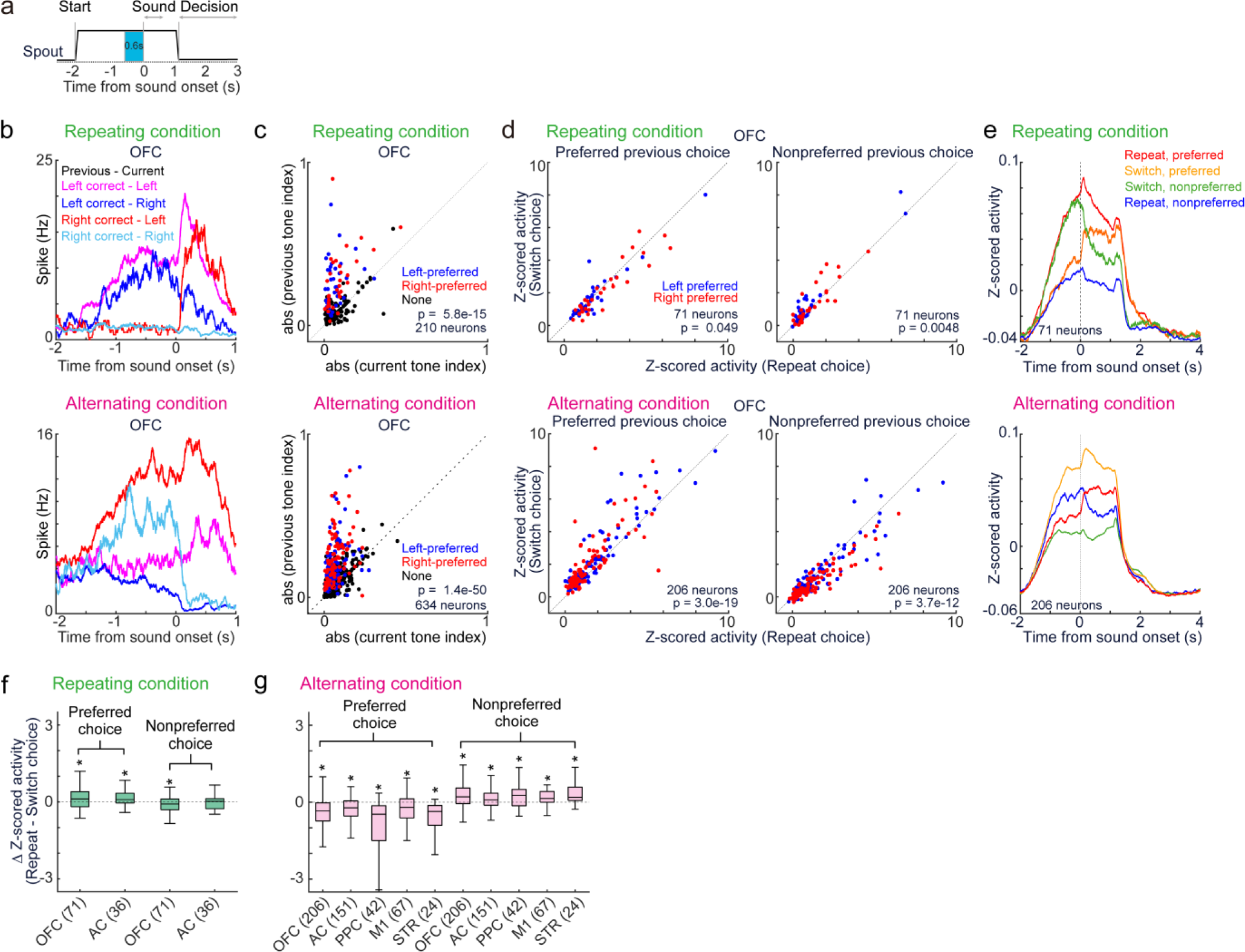
Neural activity before sound onset. **a.** Analysis of neural activity between −0.6 and 0 s from sound onset. **b.** Average activity of example neurons from OFC before sound onset. **c.** Scatterplot comparing the absolute value of tone indices for previous and current trials (**Methods**). The previous tone indices in correct trials were higher than the current tone indices, suggesting that the neurons represented previous choices (blue and red dots: left- and right-side preferred neurons, p < 0.01 in the Mann‒Whitney U test; black dots: non-side-preferred neurons; p value in the Wilcoxon signed rank test). **d.** Average activity during repeated choices (x-axis) and switched choices (y-axis) between −0.6 and 0 s from sound onset (p value in the Wilcoxon signed rank test). **e.** Average traces of neurons in the OFC. Neurons were identical to **d**. **f-g.** The difference in average activity between the repeated and switched choices before sound onset. Outliers, defined as values beyond 1.5 times the interquartile range, were excluded from the plots. The numbers in parentheses show the number of left- and right-side-preferred neurons. * p < 0.05 in the Wilcoxon signed rank test.

To confirm the neural representations of previous choices, we compared the tone indices between the previous and current trials (**Methods**). The tone index compares activity during low- and high-category sound trials and ranges between −1 and 1^31,33^. By comparing the absolute tone indices for the previous correct and current trials, we quantified whether the neurons encoded the previous or current choice (**Fig. 4c**). We found that the previous tone indices in correct trials in all the brain regions were greater than the tone indices in the current trial, suggesting that the neurons in multiple brain regions represented the previous choice (**Fig. 4c, Supplementary Fig. 4a**).

To simultaneously analyze the activity of left- and right-choice representing neurons, we defined the preferred side of each neuron based on the tone index (**Methods**). Left-side preferred neurons had tone indices less than 0 in correct trials, whereas right-side preferred neurons had tone indices greater than or equal to 0 in correct trials. These neurons had significantly different activities between the left- and right-correct choice trials (p < 0.01 according to the Mann‒Whitney U test).

We investigated whether the neural encoding of previous choices was modulated by the upcoming choice in the current trial (**Fig. 4d**). In the repeating condition, when the previous choice was the preferred choice, the OFC neurons significantly increased the activity when mice repeated the choice rather than when they switched their choice even before the sound was presented (**Fig. 4d, top left; Fig. 4e**). Conversely, in the alternating condition, the OFC neurons increased the activity when the mice switched from the previous preferred choice (**Fig. 4d, bottom left**). When the previous choice was the nonpreferred side, neurons in both the repeating and alternating conditions showed opposite activity compared to the preferred previous choice (**Fig. 4d, right**). The neurons in the AC, PPC, STR, and M1 showed activity patterns similar to those of the OFC (**Fig. 4f and g; Supplementary Fig. 4b and c**). These results suggest that the neurons in multiple brain regions not only represented the previous choice but also modulated the activity according to the upcoming choice.

### The activity of neurons during sound presentation depends on the choice sequence

During sound presentation, we identified 785 and 2535 neurons in the repeating and alternating conditions, respectively, that exhibited a significant increase in activity (21.81% and 27.79%, respectively, of the task-relevant neurons) (**Fig. 5a**). In the repeating condition, the activity of the example neuron in the OFC gradually increased when the previous trial was right-rewarded, followed by a right choice in the current trial (**Fig. 5b, top**). In the alternating condition, an example OFC neuron increased the activity when the choice was switched from left to right (**Fig. 5b, bottom**).

**Fig. 5.**
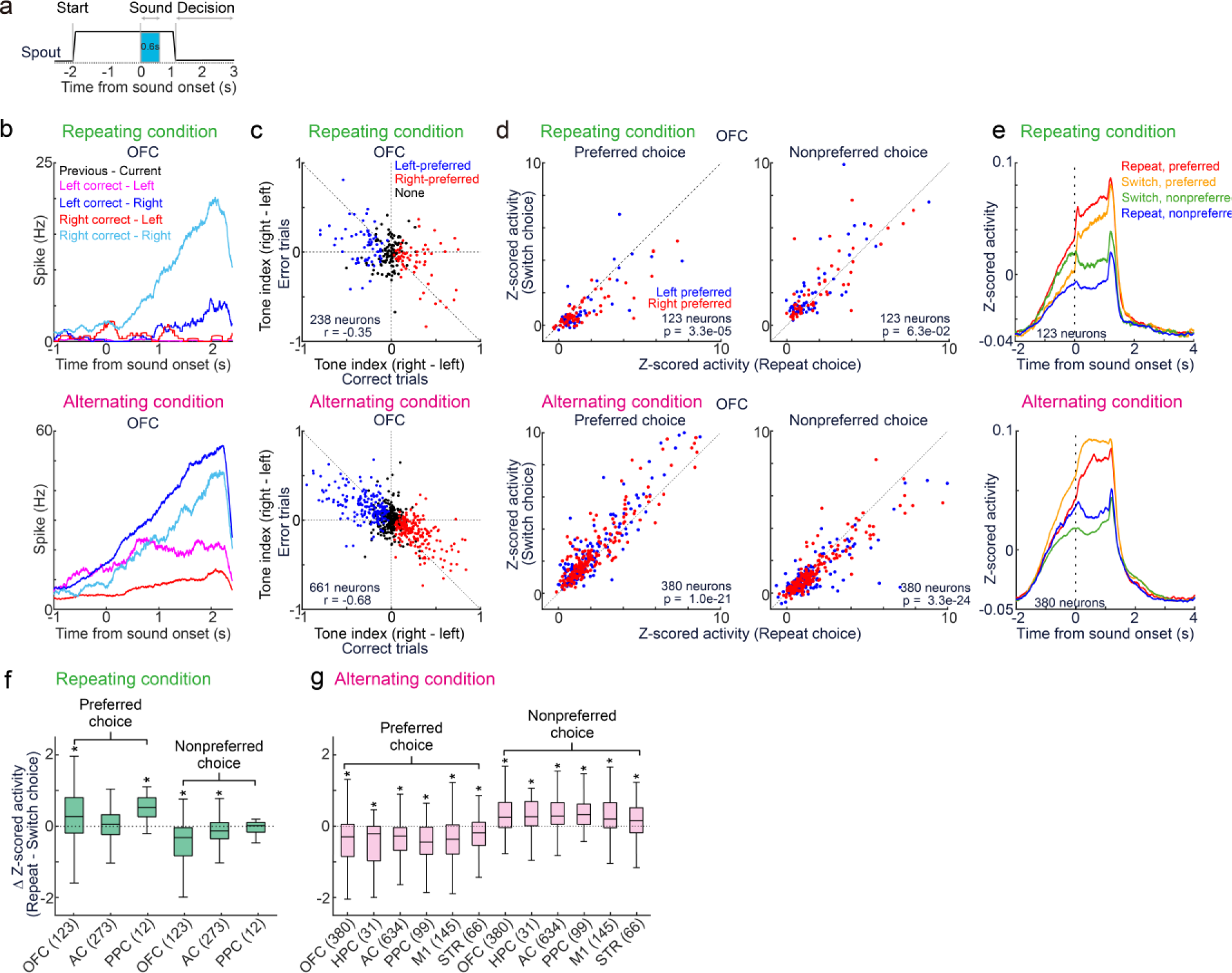
Neural activity during sound presentation. **a.** Analysis of neural activity during 0.6 s of sound. **b.** Average activity of example neurons in the OFC during sound. **c.** Tone indices of task-relevant neurons. The tone indices of OFC neurons had negative correlations between the correct and incorrect trials, suggesting choice representation. Data plots are ame as Fig. 4. **d.** Average activity of each neuron during the repeated (x-axis) and switched choices y-axis) (p value in the Wilcoxon signed rank test). **e.** Average activity of neurons in the OFC. **f-g.** The difference in average neural activity between the repeated and switched choices during sound. The parentheses show the number of left- and right-side-preferred neurons. * p < 0.05 in the Wilcoxon igned rank test.

To investigate whether the neurons represented choices or sounds, we analyzed the tone indices in the correct and incorrect trials (**Methods**). When neurons represented sound categories, the correlations of the tone indices between correct and incorrect trials were positive. Conversely, when neurons represented choices, the correlations were negative^31,33^. The tone indices of neurons in the OFC, PPC, STR, HPC, and M1 were negatively correlated between the correct and incorrect trials, suggesting the choice encoding (**Fig. 5c, Supplementary Fig. 5a**). In contrast, the tone indices of AC neurons were positively correlated with each other, suggesting that they were involved in sound encoding (**Supplementary Fig. 5a**).

We investigated whether the activity of neurons was changed by the choice sequence (**Fig. 5d**). In the repeating condition, the OFC neurons showed increased activity when the preferred choice was repeated compared with when the choice was switched to the preferred side (**Fig. 5d, top left; Fig. 5e**). In contrast, in the alternating condition, the OFC neurons increased the activity when the choice was switched to the preferred side (**Fig. 5d, bottom left**). In the nonpreferred choice condition, the OFC neurons showed opposite activity than in the preferred choice condition in both the repeating and alternating conditions (**Fig. 5d, right**). Similar to the OFC, the other recorded brain regions exhibited choice-sequence-dependent activity (**Fig. 5f and g; Supplementary Fig. 5b and c**). Given that repeated and switched choices had high-reward expectations in the repeating and alternating conditions, respectively, these results suggest that the neurons in various brain regions encode reward expectations in both model-free and inference-based strategies.

### Global neural encoding of unexpected outcomes in the repeating but not in the alternating condition

At the time of the outcome (**Fig. 6a**), we identified 1895 and 4491 task-relevant neurons in the repeating and alternating conditions, respectively (52.56% and 49.08% of the task-relevant neurons). An example OFC neuron in the repeating condition showed increased activity when the mouse switched its choice from right to left rather than when it repeated the left choice (**Fig. 6b, top**). In contrast, in the alternating condition, the OFC neuron did not show choice sequence-dependent activity (**Fig. 6b, bottom**). The tone index during the outcome timing suggested that all the recorded regions had choice encoding (**Fig. 6c; Supplementary Fig. 6a**).

**Fig. 6.**
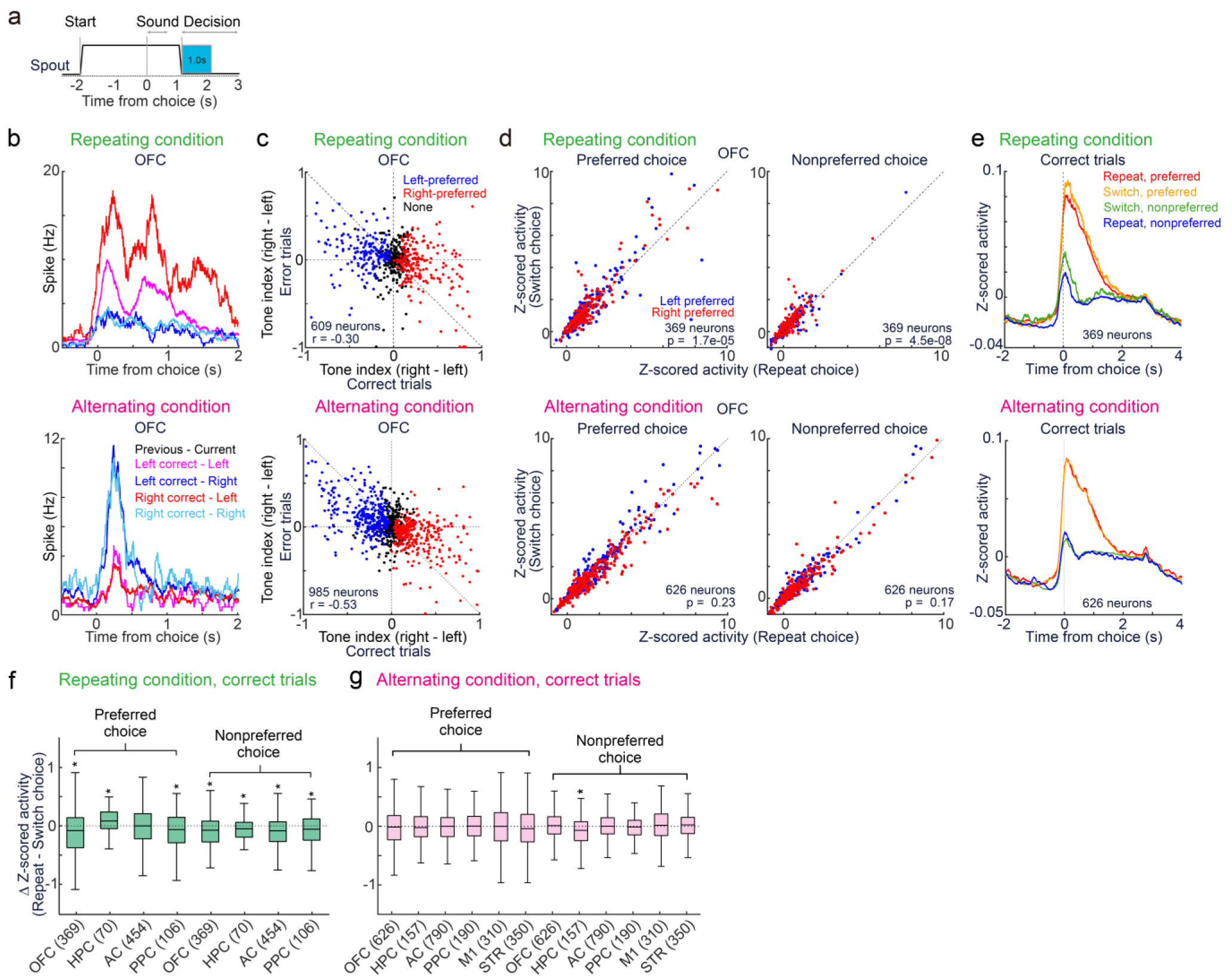
Neural activity at outcome timing. **a.** Analysis of neural activity between 0 and 1.0 s from choice and outcome. Data presentations are consistent with those in Figs. 4 and 5. **b.** Average activity of example neurons during the outcome including both correct and incorrect trials. **c.** Tone indices of task-relevant neurons. The tone indices of OFC neurons suggested the choice representation (blue, red, and black dots: left-, right-, and non-side-preferred neurons, respectively). **d.** Comparison of activity between the repeated and switched choices (p value in the Wilcoxon signed rank test). **e.** Average activity of neurons in the OFC. **f-g.** The difference in average activity between the repeated and switched choices in the current correct trials. Outliers are not shown in the plots. The parentheses show the number of left- and right-side-preferred neurons. * p < 0.05 in the Wilcoxon signed rank test.

We analyzed whether the change in activity of the neurons depended on the previous choice (**Fig. 6d**). In the correct trials in the repeating condition, the OFC neurons exhibited increased activity for the preferred choice when the choice was switched compared with when it was repeated (**Fig. 6d top left, 6e, 6f**). In contrast, in the incorrect trials, OFC neurons exhibited greater activity when choices were repeated than when choices were switched (**Supplementary Fig. 7**). Given that reward expectations were lower in switched choices than in repeated choices, these results suggest that the neurons in the OFC encode unexpected outcomes. We found similar activity patterns in the PPC, HPC, and AC (**Supplementary Figs. 6 and 7**), suggesting that multiple brain regions globally represented unexpected outcomes for the model-free strategy.

In contrast, in the alternating condition, none of the recorded brain regions showed choice sequence-dependent activity in the correct trials (**Fig. 6g, Supplementary Fig. 6**). In the incorrect trials, only the PPC neurons exhibited a change in activity based on the reward expectation (**Supplementary Fig. 7**). During the overtrained phase of electrophysiological neural recording, the mice exhibited experience-dependent choice updates in the repeating condition, while their behavior was stable in the alternating condition (**Fig. 2g and h**). These results were consistent with the activity at the outcome time, in which the activity was modulated by the reward expectation mainly in the repeating condition.

## Discussion

To investigate the neural representation of model-free and inference-based strategies in multiple brain regions, we used a tone frequency discrimination task in head-fixed mice with different transition probabilities in the tone category. We found that mice tended to repeat previously rewarded choices even when the tone category was randomly selected in the neutral condition (**Fig. 1c**). This default strategy was continuously used in the repeating condition to properly bias the choices. In contrast, mice took several sessions to reverse the choice biases in the alternating condition (**Fig. 1e**).

An RL model estimating the expected outcome of each choice fit the mice’s choices in the neutral and repeating conditions, suggesting that the default model-free strategy was used in the repeating condition. In the alternating condition, the behavioral strategy gradually shifted from the model-free to the state-based model, which determined a hidden context by estimating the transition probability of tone category (**Fig. 2**). During the overtrained phase with an electrophysiological recording with Neuropixels, the behavior of the mice was fit to the model-free and inference-based strategies under the repeating and alternating conditions, respectively (**Fig. 2i**).

In both conditions, the neurons in all the recorded regions, including the OFC, PPC, HPC, STR, M1, and AC exhibited increased activity when the preferred choices or tones expected a large reward probability, suggesting that brain-wide encoding of reward expectations occurred both in the model-free and inference strategies (**Figs. 4–5**). In contrast, at the time of the outcome, the neurons exhibited increased activity with unexpected outcomes only in the repeating condition (**Fig. 6**). This was consistent with the behavioral data: during the overtrained electrophysiological phase, the choices of the mice were stable under the alternating condition, while the choices were updated by past experiences in the repeating condition (**Fig. 2h**). Our results suggest the global neural encoding of reward expectations in both the model-free and inference-based strategies.

The behavioral strategy of repeating a previously rewarded choice, such as the ‘win-stay lose-switch strategy’, has been observed in previous studies with humans^34^, primates^35^, and rodents^29^. We found that mice tended to repeat rewarded choices even when the sound stimuli were randomly presented in the neural condition (**Fig. 1c**). This default strategy was continued to optimize choices in the repeating condition, resulting in the rapid acquisition of proper choice biases. In contrast, mice required several sessions to achieve alternating choice biases in the alternating condition (**Fig. 1e and Supplementary Fig. 1**). A previous study involving a probabilistic foraging task showed that mice used a stimulus-bound model-free strategy in the early phase of training and gradually shifted this strategy to an inference-based strategy^8^. This was consistent with our results in the alternating condition in which the state-based model gradually fit the mouse behavior better than model-free RL (**Fig. 2e**). Similar to our behavioral task, previous studies in free-moving rodents used a perceptual decision-making task with a transition probability in the sensory category^8,28,29^. These studies trained one animal both in the repeating and alternating conditions and observed the use of a task-specific inference-based strategy in both conditions. In contrast, we trained separate head-fixed mice in either the repeating or alternating condition and found differences in learning speed and strategy between the two task conditions (**Fig. 1e and 2**).

The electrophysiological recordings showed that the choices and outcomes were globally represented in the cortical and subcortical regions, while the sound stimuli were selectively represented in the auditory cortex during sound presentation, irrespective of the task condition (**Fig. 3d and e**). Interestingly, the AC neurons represented choices before and after the sound presentation, consistent with previous findings^16^.

We also found that the modulation of neural activity based on reward expectations was globally observed in the cortical and subcortical regions for both the model-free and inference-based strategies (**Figs. 4–6**). Previous studies have shown that the OFC plays an essential role in both model-free^36^ and inference-based flexible behavior^8^. The AC neurons encode task rules and reward expectations^16,37,38^. The HPC, PPC, and STR are involved in either the model-free or model-based strategy^24,39,40^. These studies led to the hypothesis that behavioral strategies are globally encoded in the brain^30^. However, as many studies have targeted a specific brain region for a specific behavioral strategy, it was unclear whether the neural representations of model-free and inference-based strategies were widely distributed in the brain. Our study suggested that, at least in the overtrained phase, the reward expectations of both the model-free and inference-based strategies were globally represented in the brain.

At the outcome timing, in the repeating condition, we found that the neural activity of OFC, PPC, HPC, and AC increased when the mice switched the choice and rewarded (**Fig. 6**). Switching behavior was uncommon in the repeating condition; thus, reward expectations for switching were lower than those for repetition. These neural representations of unexpected outcomes are important for computing a reward prediction error for value updating in the model-free strategy^5,41,42^.

In the alternating condition, we did not observe neural encoding of unexpected outcomes, possibly because of stable mouse behavior (**Fig. 2h**). Additional studies are required to investigate how the inference-based strategy is computed in the brain. Additionally, it is important to investigate neural representation during the learning phase to determine how model-free and inference-based strategies are acquired in the brain.

In summary, we found that mice used the default model-free strategy to bias choice in both the neutral and repeating conditions, while the choice behavior of mice changed to an inference-based strategy in the alternating condition. In the overtrained phase, the neural activity of the frontal and sensory cortices, hippocampus, and striatum was correlated with the reward expectation of both the model-free and inference-based strategies. Neurons in multiple brain regions exhibited increased activity with unexpected outcomes in the repeating condition. These results propose the global encoding of model-free and inference-based strategies in the brain.

## Materials and Methods

All animal procedures were approved by the Animal Care and Use Committee at the Institute for Quantitative Biosciences (IQB), University of Tokyo. Mice were housed in a temperature-controlled room with a 12 h/12 h light/dark cycle. All the experiments were performed during the dark cycle.

### Subjects

Male CBA/J mice (n = 26, Strain #000656; The Jackson Laboratory), aged 8 to 15 weeks at the start of behavioral training, were used for the experiments. 26 mice were allocated for behavioral experiments. 14 out of the 26 mice were subjected to electrophysiological recording after the behavioral experiments. Before surgery, 3 mice were housed in one cage. Mice were allowed ad libitum access to food, while water intake was restricted to 1.5 mL per day. On weekends, the mice were given 3 mL of extra water and free access to 1.5% citric acid water to prevent dehydration. Mice were caged in isolation after craniotomy.

### Surgery

The surgical procedures were described in our previous research^1,31^. In summary, the surgery had two steps. First, a custom-designed head bar was implanted for behavioral training. Second, a craniotomy was performed for electrophysiological recording.

For head bar implantation, the mice were anesthetized via intraperitoneal injection of a mixture of medetomidine (0.3 mg/kg), midazolam (4.0 mg/kg), and butorphanol (5.0 mg/kg). Meloxicam (2.5 mg/kg) and eye ointment were also used. Mice were placed in a stereotaxic apparatus. The scalp was removed above the entire cortical area. We cleaned the skull with povidone iodine and hydrogen peroxide. We attached the head bar to the skull with Superbond adhesive (Sun Medical or Parkell S380) and cyanoacrylate glue (Zap-A-Gap, PT03)^43^.

For craniotomy, the mice were anesthetized with isoflurane (2% for induction, 1.5% for maintenance). We targeted 6 brain regions: the OFC, STR, M1, PPC, HPC, and AC. The OFC sites were +2.6 mm anterior-posterior (AP) and ±1.4 mm medio-lateral (ML) from bregma. The STR and M1 sites were +0.8 mm AP and ±1.5 mm ML. The PPC and HPC sites were −2.0 mm AP and ±1.7 mm ML. The AC sites were −3.0 mm AP and ±3.8 mm ML (**Supplementary Fig. 2**). We drilled a small hole (0.5−0.8 mm in diameter) through cyanoacrylate glue and the skull to expose the brain surface and removed the dura mater. We covered the brain surface with agar dissolved in PBS followed by silicone oil and Kwik-Sil (World Precision Instruments) to prevent drying.

### Behavioral apparatus

We performed behavioral experiments inside a custom-made training box or a sound-attenuating booth (O’hara, Inc.). After recovering from the head bar implantation, the mice were head-fixed and placed on a custom cylinder treadmill. We presented sound stimuli precalibrated with a Brüel and Kjaer microphone (Type 4939) from an Avisoft Bioacoustics speaker (#60108) positioned to the right front of the mice. Two spouts were placed in front of the mice to deliver sucrose water. The licking behavior of the mice was detected using electrical or infrared sensors^25^. We used custom-made MATLAB (MathWorks) programs on Bpod r0.5 (https://sanworks.io) on Windows OS.

### Tone frequency discrimination task with probabilistic alternation of tone category with transition probabilities

Like in our previous studies^1,31^, each trial began by retracting the two spouts away from the mice. After a random interval of 1 to 2 seconds, a sound stimulus in the form of a tone cloud was presented^25–27^. The intensity of the tone cloud in each trial remained constant but was sampled from 60, 65, or 70 dB SPL (the sound pressure level in decibels with respect to 20 µPa). The duration of the tone cloud was held constant at 0.6 s. The tone cloud was a mixture of low-frequency (5–10 kHz) and high-frequency (20–40 kHz) tones.

After the sound ended, the two spouts were immediately displayed to the mice. Mice selected either the left or right spout depending on the dominant tone frequency. The association between the tone category and rewarded choice was determined for each mouse. A correct choice provided 2.4 µL of 10% sucrose water. An incorrect choice triggered a 0.2-s noise burst from the speaker. If the mouse failed to select a spout within 15 seconds, a new trial started.

Our behavioral task had 4 steps. 1) The initial step involved training the mice to discriminate between the 90% low-frequency and 90% high-frequency tone clouds. We trained 26 mice in the initial step. 2) After the mice were able to discriminate the sounds, the neutral condition started. We used 6 tone clouds (0, 20, 35, 65, 80, and 100% high-frequency tones) with presentation probabilities of 25%, 12.5%, 12.5%, 12.5%, 12.5%, and 25%, respectively. The neutral condition had a transition probability ‘*p*’, which controlled how often the tone category of the previous trial alternated in the current trial. We set ‘*p* = 0.5’ to randomly present the tone category in each trial. 7 out of the 26 mice skipped step 2 and directly completed step 3.

3) After the mouse experienced at least one training session in the neutral condition or when the percentages of correct responses for tone clouds that were 90% high tones and 90% low tones were both greater than 80% in the initial step, we assigned the mouse to either the repeating or alternating condition (22 out of 26 mice). In both conditions, we used 6 tone clouds (0, 25, 45, 55, 75, and 100% high-frequency tones) with presentation probabilities of 25%, 12.5%, 12.5%, 12.5%, 12.5%, and 25%, respectively. The repeating condition had a transition probability of ‘*p* = 0.2’, in which the same tone category was frequently presented. In contrast, in the alternating condition, we set the transition probability as ‘*p* = 0.9’, where the tone category was alternated every trial in 90% of trials. In both the repeating and alternating conditions, each session started with 40 trials of only 100% low- or high-tone clouds. The repeating condition switched the tone category in every 10 trials, while the alternating condition switched the category every trial. After the first 40 trials, the above conditions for the 6 tone clouds and transition probabilities started. 3 mice did not show transition probability-dependent choice biases, as analyzed with a psychometric function (**Methods, Behavioral analysis**)^1^, in the alternating condition after more than 11 sessions and were not used for the analyses.

4) After the mice experienced at least 9 sessions of either the repeating or alternating condition, we started the prerecording step. This step gradually increased the interval between the end of the sound and the spout approaching until the interval reached 0.5 s. The transition probability of the alternating condition was set to ‘*p* = 0.8’. 14 mice completed the prerecording step.

### Electrophysiological recording and histology

The electrophysiological recording performed with Neuropixels 1.0 (IMEC) and the histological analysis were described in detail in our previous study^31^. In summary, we inserted the Neuropixels probe 1 to 3 times in each hole in both the left and right hemispheres of the brain. For the OFC recordings, the probe was tilted 5 degrees in the medial direction. For the AC recordings, the probe was tilted 18 degrees in the lateral direction. The angle was 0 degrees for the PPC, HPC, STR, and M1 recordings. To identify the probe location in the post hoc fixed brain, the probe was soaked in a diluted solution of CM-DiA or DiI (Thermo Fisher Product #D3883 or #V22888). The probe was manually lowered to the brain surface through a mixture of agar and PBS at a speed of 120 μm/min (MPC−200 Controller and ROE, Sutter Instrument)^44^. The brain surface was defined based on the depth at which spikes were initially observed at the recording electrodes. The 384 electrodes from the tip of the Neuropixels probe were used for recording. The Open-Ephys GUI acquired the neural data at a sampling rate of 30 kHz with a gain of 500 (PXIe acquisition module, IMEC). The task events, including treadmill rotations, were sampled at 2.5 kHz (BNC−2110, National Instruments). After the recording was complete, the probe was slowly extracted, and the hole was covered with Kwik-Sil.

After the electrophysiological recording, the mice were deeply anesthetized with isoflurane (5%) and further anesthetized with a mixture of 1.5 mg/kg medetomidine, 20 mg/kg midazolam, and 25 mg/kg butorphanol. Mice were perfused with 10% formalin solution. Brain sections were sliced with a vibratome to a thickness of 100 µm (VT1000S; Leica Biosystems) and mounted with DAPI mounting medium (Vector Laboratories, Cat. No. H−1200). The probe locations were captured with a confocal laser scanning microscope (FV3000, Olympus) at 4x magnification (**Fig. 3**).

### Data analysis

#### Number of mice and sessions in behavioral tasks

We used the sessions for analyses for which (i) the correct response rate of the mouse for both the 100% low and 100% high tones exceeded 75% and (ii) the total reward amount in one session was above 600 µL. Under neutral conditions, we analyzed 28 sessions from 16 out of 19 mice (**Fig. 1**); the sessions in 3 mice did not exceed the accuracy rate of 75%. Under the repeating and alternating conditions, we analyzed 113 and 141 sessions, respectively, from 8 and 11 mice (**Fig. 1**). The behavioral results of (i) all the sessions (neutral condition: 35 sessions in 19 mice; repeating and alternating conditions: 139 and 165 sessions from 8 and 11 mice) and (ii) the sessions with correct rates over 80% (108 and 117 sessions from 8 and 10 mice) are shown in **Supplementary Fig. 1**.

#### Behavioral analysis

We analyzed the choice behavior of mice with a psychometric function based on our previous study^1^. The psychometric function models the perceptual uncertainty of mice with a truncated Gaussian ranging between 0 and 1^45^. Our model investigated whether the choice biases of mice depended on the rewarded side in the previous trial. We tested whether the psychometric function with a choice-bias parameter better fit the mouse choices than that without the parameter (p < 0.01 in the likelihood ratio test). We investigated when the mouse started to bias the choices in the repeating and alternating conditions (**Fig. 1e**).

#### Behavioral model

The behavioral models were based on signal detection theory (SDT). The behavioral task required the mice to estimate the hidden state (*S*) of left-rewarded (*S* = *L*) or right-rewarded (*S* = *R*) based on the sensory evidence of the tone cloud. SDT shows that both the expected outcome in each state and the belief state probability are essential for optimizing choices^1,2^. The following model-free reinforcement learning (RL) model and state-based model estimated the expected outcome and state probability, respectively.

##### Model-free reinforcement learning (RL) model

The RL model updated the expected outcome of left and right choice in each state *Q*_*a*,*S*_, defined as the prior value^1,31,46^, while the belief state probability was fixed. We denoted the choice as *a*. We assumed that there were only left-rewarded and right-rewarded states; the prior values satisfied the criteria *Q*_*left*,*R*_ = *Q*_*right*,*L*_ = 0. We simplified *Q*_*left*,*L*_ and *Q*_*right*,*R*_ as *Q*_*left*_ and *Q*_*right*_, respectively. The model used prior values to compute the decision threshold *x0* in each trial (*t*) with a softmax function and an inverse temperature parameter *β*:

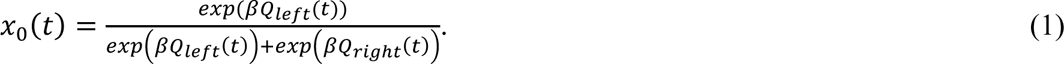

The softmax equation modeled a perceived reward size that might be different from the actual amount of water. The right-choice probability at trial *t*, *P*(*right*, *t*), was estimated from a perceptual uncertainty *σ* and a bias parameter *d*:

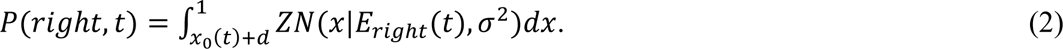

*Eright*(*t*) was the proportion of tone frequency associated with a rightward choice in a tone cloud. *Z* truncated the Gaussian distribution between 0 and 1, here and hereafter. We updated the prior value with forgetting Q-learning^15^:

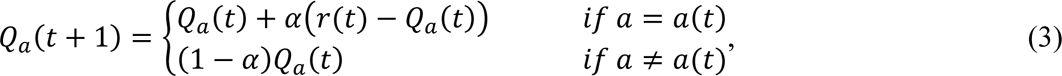

where *α* was the learning rate. *r*(*t*) was the outcome at trial *t*. The initial prior value for each choice was the amount of reward (i.e., 2.4).

##### State-based model

The state-based model had the belief of state probability *P*(*S*) in each trial by estimating and updating the transition of state *P*_*transition*_ in every trial^7^. The prior values were fixed. Bayesian inference provided the decision threshold of choice based on *P*(*S*). First, the likelihood of a sensory stimulus *x* in state *Si*, *P*(*x*|*Si*), was defined as follows:

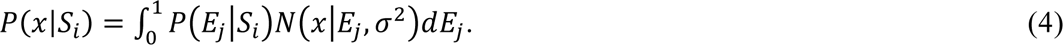

*P*(*E*_*j*_|*S*_*i*_) was the probability of tone cloud *Ej* in a given state *Si*. *σ* was a free parameter for perceptual uncertainty. The posterior probability *P*(*Si*|*x*) was calculated with the Bayes rule:

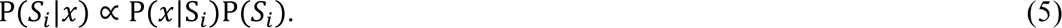

The decision threshold *x*_0_ satisfied P(*S*_*L*_|*x*_0_) = P(*S*_*R*_|*x*_0_). We used a softmax equation to add flexibility to the threshold *x*_0_ with an inverse temperature parameter *β*:

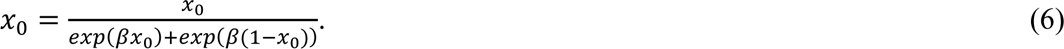

Based on *x*_0_ and the bias choice parameter *d*, the model estimated the choice in each trial with Equation (2).

After the model received the outcome at trial *t*, the state transition *P*_*transiton*_ was updated based on the true state of trial *t* and *t*−1:

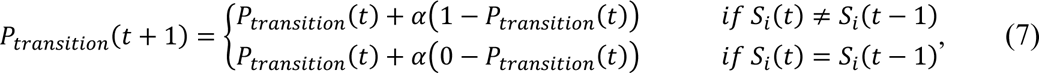

where *α* was the learning rate. The model computed the belief state of trial *t*+1 based on the transition probability and the true state at *t*. The initial prior belief of left- and right-rewarded states was 0.5. The initial transition probability was 0.5 for the learning phase of the repeating and alternating conditions, while it was the true transition probability for the overtrained phase.

### Model comparison

We defined the likelihood *l*(*t*) from the estimated choice probability in each trial with Equation (2):

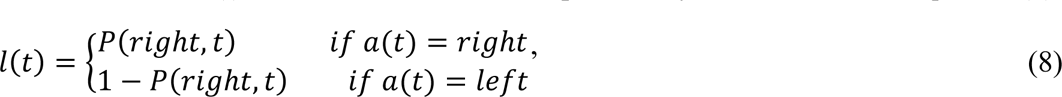

We then analyzed the likelihood in each session *L* using the trials without the first 40 trials in each session:

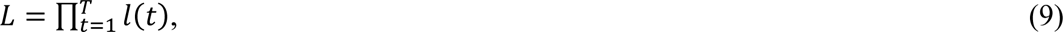

where *T* was the number of trials. The model parameters were fit to achieve the maximum likelihood. We first used the Bayesian information criterion (BIC) to identify the necessary parameters in the RL model or state-based model. We also used the BIC to compare the performance of the RL and state-based models (**Fig. 2**):

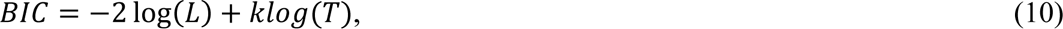

where *k* was the number of free parameters.

### Number of mice and sessions of electrophysiological neural recording

Same as the behavioral data analyses, we used the sessions for analyses when (i) the percentage of correct responses for both the 100% low tones and 100% high tones stimuli were greater than 75% and (ii) the total reward amount in one session was at least 600 µL. We analyzed OFC neurons from 17 and 30 sessions from 5 and 7 mice in the repeating and alternating conditions, respectively; PPC and HPC neurons from 18 and 39 sessions from 4 and 9 mice; and AC neurons from 16 and 31 sessions from 4 and 8 mice. The STR and M1 neurons from 39 sessions from 7 mice were analyzed only in the alternating condition. 13 and 36 sessions in the repeating and alternating conditions, respectively, were excluded from the analyses.

### Electrophysiology data analysis

Spike sorting and manual curation were performed with KiloSort−3 on MATLAB (https://github.com/cortex-lab/KiloSort) and Phy on Python (https://github.com/cortex-lab/phy). KiloSort−3 spike sorting tracked the approximate depth of spikes from each unit during a session. We defined the depth of each unit from the approximate location of the probe and the electrode position, which measured the maximum amplitude of spikes on average.

For a Neuropixels probe for the OFC, we used the units for analyses when the estimated spike depth from the brain surface was less than 1.9 mm. For the PPC recording probe, we used units for analyses when the estimated spike depth was less than 1.0 mm. For the STR recording probe, we analyzed the units when the estimated spike depth was greater than 1.5 mm. For the M1 probe, we used units when the estimated spike depth was less than 1.5 mm. For the HPC, we used the units for analyses when the estimated spike depth was between 1 and 2.3 mm. We analyzed all the units recorded by the AC probe.

We identified task-relevant neurons that exhibited increased activity during the task (p < 1.0e−10 in the one-sided Wilcoxon signed rank test) compared to baseline activity. For sound-aligned activity, task-relevant neurons exhibited increased activity in at least one time window (0.1 s) between −1.5 s and 2.5 s from sound onset (40 windows). For choice-aligned activity, task-relevant neurons exhibited increased activity in at least one time window between −0.5 and 2.5 s from the choice (30 windows, in total 40 + 30 = 70 windows). The baseline for the sound-aligned activity was −0.2 to 0 s from spout removal, i.e., before trial initiation. The baseline for the choice-aligned activity was −0.2 to 0 s from the time the spout approached, i.e., between the end of the sound and making a choice.

Among the task-relevant neurons, we focused on the neurons that exhibited significantly increased activity (1) between −0.6 and 0 s from sound onset (before sound), (2) between 0 and 0.6 s from sound onset (during sound), and (3) between 0 and 1.0 s from choice (during outcome).

We analyzed the tone index of each neuron based on the high- and low-category tone trials:

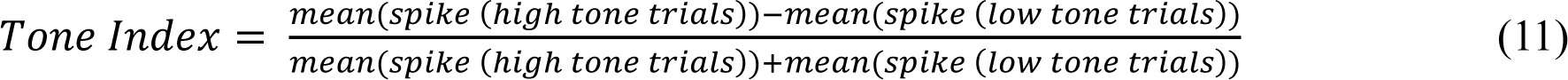

The tone index ranged between −1 and 1 and was independently analyzed for correct and incorrect trials (**Figs. 5 and 6**). In the analyses of neural activity before sound presentation (**Fig. 4**), we analyzed the tone indices of the previous and current trials to determine whether the neurons represented previous or current choices. With respect to the tone indices in previous correct trials, we independently analyzed the tone indices in the current left- or right-choice trials and averaged the values (previous tone indices). For the current tone index, we independently analyzed the tone index in the previous left or right correct choice trials and averaged the values.

We defined the preferred side of the task-relevant neurons based on (i) the tone index in the correct trials and (ii) the activity difference between the left and right correct choice trials. First, left- and right-side preferred neurons had negative and positive tone indices, respectively, in the correct trials. Second, the activity on the preferred side was greater than that on the nonpreferred side (Mann‒Whitney U test, p < 0.01). For example, when the tone index of a neuron was less than 0 and the activity in the correct left-choice trials was greater than that in the correct right-choice trials, the preferred side was defined as the left side (**Supplementary Fig. 3c**). If neurons did not show a significant difference in activity between the left and right correct choice trials (p > 0.01), we defined them as non-side-preferred neurons (**Figs. 4–6; Supplementary Figs.4–6**).

### Regression analysis

A generalized linear model was used to analyze whether the activity of a specific time window in the task represented the choice (*C*), sound (*S*), or outcome (*O*) in both the current and previous trials, in addition to the running speed (*R*) (MATLAB, glmfit with Poisson distribution, p < 0.01) (**Fig. 3c**):

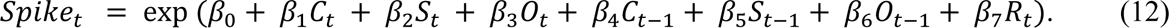

*Spike_t_* represented the number of spikes in the time windows before sound presentation, during the sound, or during the outcome. *β*_0−7_ were the regression coefficients.

### Quantification and statistical analysis

We used MATLAB 2022b for all the analyses except for spike sorting, which was performed with KiloSort 3 on Python. The statistical details are shown in the Results section, the figures, and the figure legends. Solid lines and shaded areas represent the means and standard deviations or standard errors, respectively. In the analyses of psychometric function, we used the likelihood ratio test to investigate whether the additional parameter of choice bias significantly improved the fit to mouse choices (**Fig. 1e**). Model fitting of the RL and state-based models was performed with the Bayesian information criterion (BIC) (**Fig. 2**). For the behavioral analyses of multiple sessions in each mouse, we employed the linear mixed-effects model (MATLAB: fitlme). In other analyses, we used two-sided nonparametric statistical tests. We used the MATLAB glmfit function to analyze the proportion of neurons representing each task variable (**Fig. 3d and e**). We used the chi-square test to compare the proportion of neurons representing task variables.

## Acknowledgments

This work was funded by JSPS Kakanhi (JP21H05243, JP 21H03492, JP 22H04766), AMED JP23wm0525008, and the Uehara Memorial Foundation for A.F.

## Author contributions

S.W. collected and analyzed the data and wrote the paper. H.G. and K.I. analyzed the data. A.F. designed the experiment, analyzed the data, and wrote the paper.

## Materials and Correspondence

Should be addressed to A.F.

---

**Supplementary Fig. 1.**
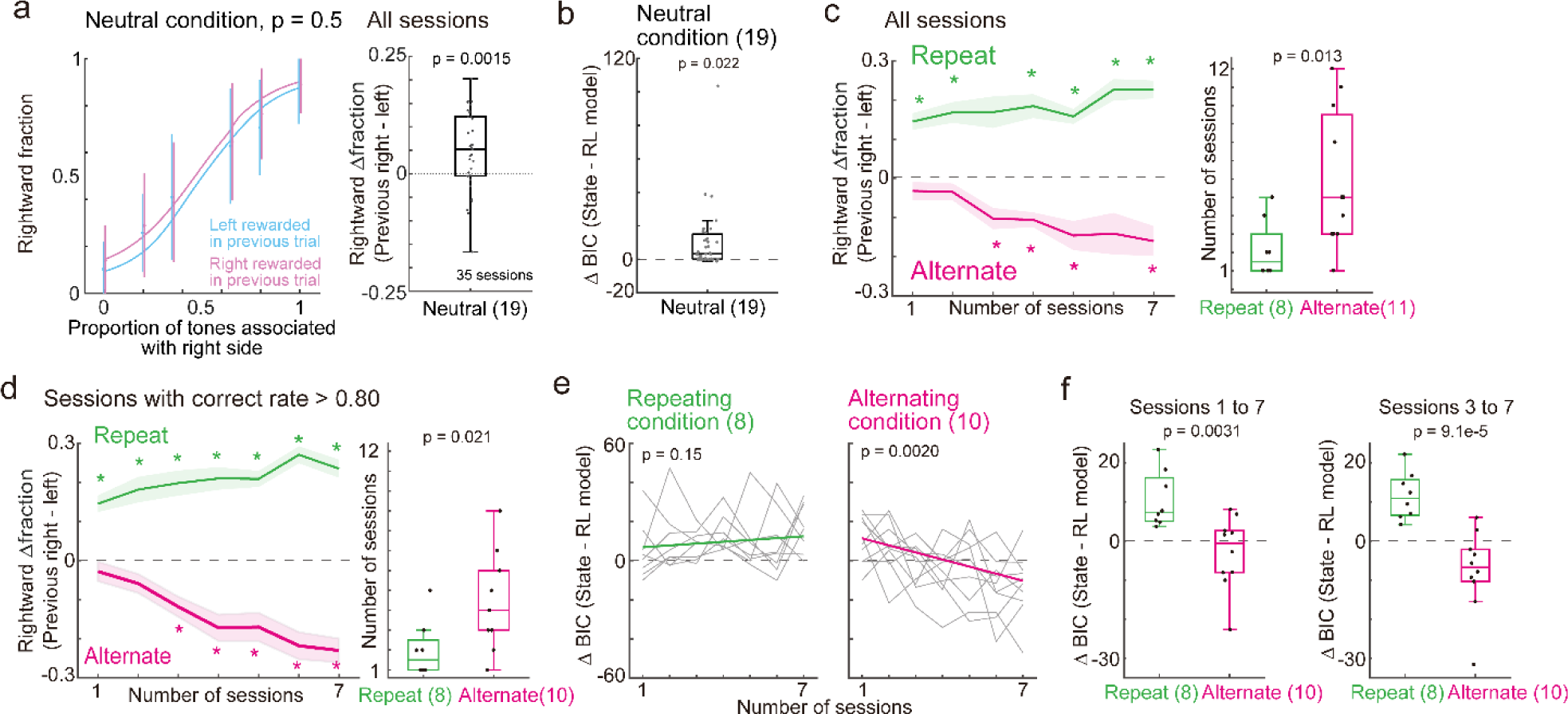
Choice behavior in all sessions and sessions with a correct response rate above 80%. **a.** Means and standard deviations of psychometric functions in the neutral condition across sessions (transition probability *p* = 0.5). Left: Data are presented in the same way as in Fig. 1c but with all the sessions, including those with a correct response rate less than 75% (35 sessions in 19 mice). Mice showed a greater probability of choosing the right side after being rewarded on the right side than after being rewarded on the left side in previous trials. Right: difference in the average fraction of right-side choices after right-side and left-side rewarded trials (p value in the linear mixed-effects model; median: central mark in the box; edges of the box: first quartile (Q1) and third quartile (Q3); bars: most extreme data points without outliers; here and hereafter). **b**. Same as Fig. 2d but for all the sessions (35 sessions in 19 mice) (p values in the linear mixed effects model). The box plot represents the difference in model fitting between the state-based and RL models. Δ BIC: Δ Bayesian information criterion. **c.** Same as Fig. 1e but for all the sessions (3 mice that did not show biased behavior in the alternating condition were excluded (**Methods**)). Left: 56 and 77 sessions from 8 and 11 mice in the repeating and alternating conditions, respectively. * p < 0.01 in the Wilcoxon signed rank test. Right, p value of the Mann‒Whitney U test. **d.** Same as Fig. 1e but with sessions in which the correct rates for easy tone clouds were above 80%. Left: 56 and 70 sessions from 8 and 10 mice in the repeating and alternating conditions, respectively; 1 mouse in the alternating condition did not reach a correct response rate of 80%. * p < 0.01 in the Wilcoxon signed rank test. Right, p value of the Mann‒Whitney U test. **e, f**: Same as Fig. 2e and 2f but for sessions in which the correct response rate in the easy-tone cloud was above 80%. **e.** We analyzed whether the regression coefficients for slope were significantly positive or negative (p value from the Wilcoxon signed rank test). The numbers in parentheses indicate the number of mice. **f**. P value of the Mann‒Whitney U test (8 and 10 mice).

**Supplementary Fig. 2.**
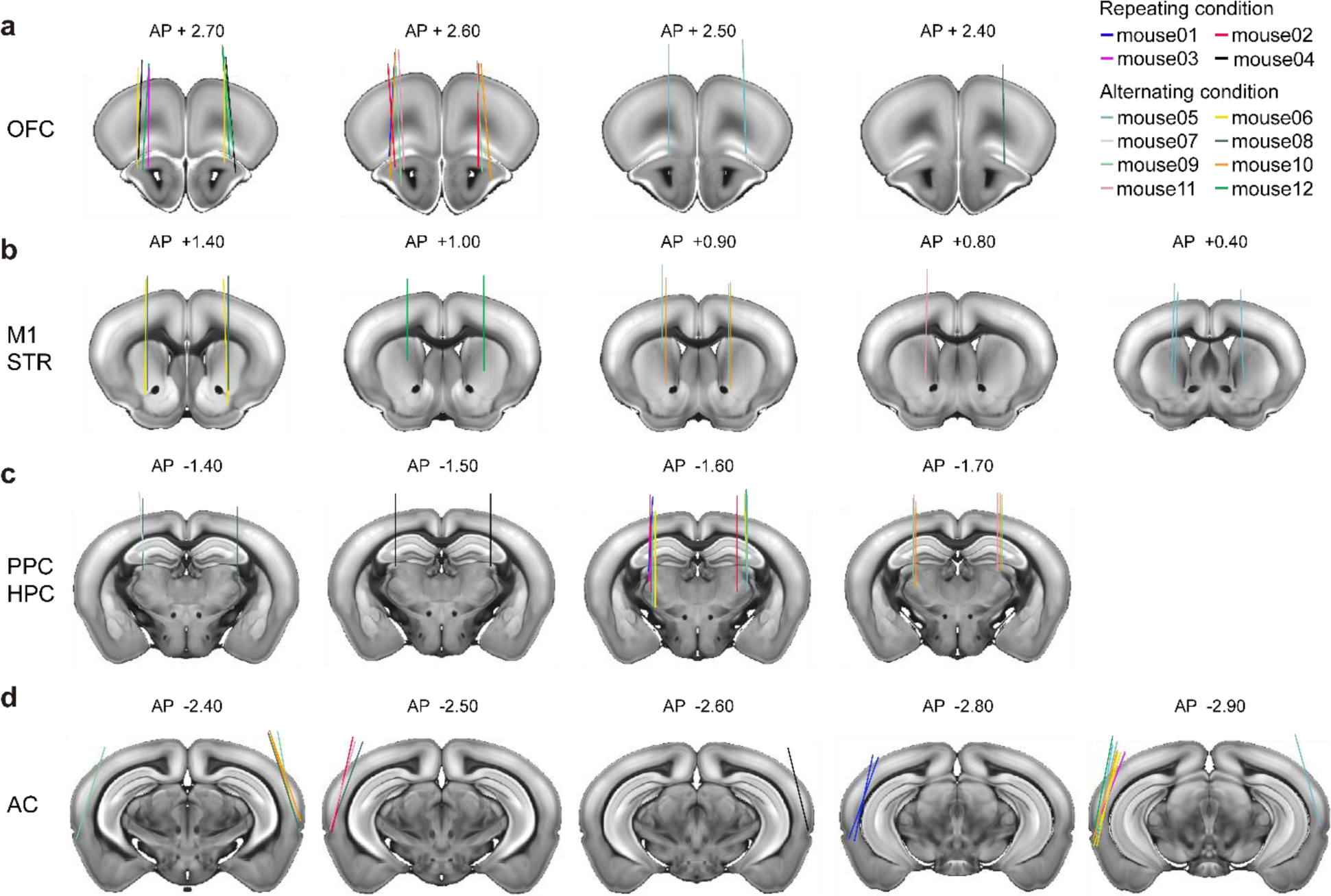
Locations of the Neuropixels probes. Probe locations were identified based on the Allen Brain Atlas coordinates. **a** to **d**: OFC, M1 and STR, PPC and HPC, and AC. Probe traces in 12 mice on coronal brain slices were registered using a Neuropixels trajectory explorer (https://github.com/petersaj/neuropixels_trajectory_explore). AP indicates the anterior-posterior axis distance from bregma (mm). In each brain region and hemisphere, we prepared a hole with a diameter of 0.5−0.8 mm in the skull for probe insertion. We identified probe locations in 17 out of the 20 holes (85%) in the OFC, 11 out of the 12 holes (91.67%) in the M1 and STR, 19 out of the 23 holes (82.61%) in the PPC and HPC, and 16 out of the 21 holes (76.19%) in the AC. All the identified probe locations were correctly placed in the target regions.

**Supplementary Fig. 3.**
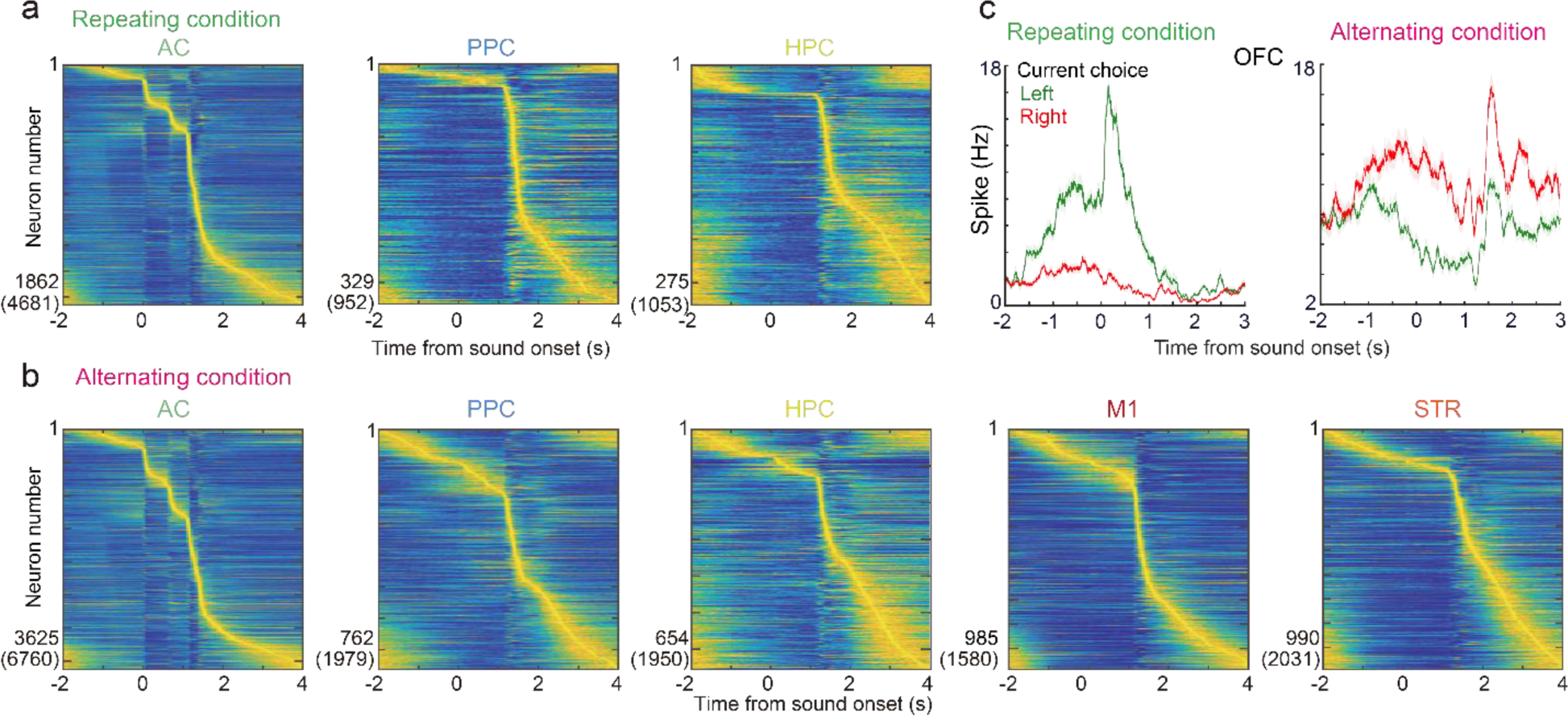
Task-relevant neurons in the auditory cortex (AC), posterior parietal cortex (PPC), hippocampus (HPC), primary motor cortex (M1), and striatum (STR). **a, b.** Average activity of task-relevant neurons (p < 1.0e−10 in the Wilcoxon signed rank test) in the repeating (**a**) and alternating conditions (**b**). The activity of each task-relevant neuron was aligned by the maximum activity timing and normalized between 0 and 1. The y-axis shows the number of task-relevant neurons. The parentheses show the number of all the identified neurons. **c.** Example left- and right-side preferred neurons in the OFC in the repeating and alternating conditions. Means and standard errors.

**Supplementary Fig. 4.**
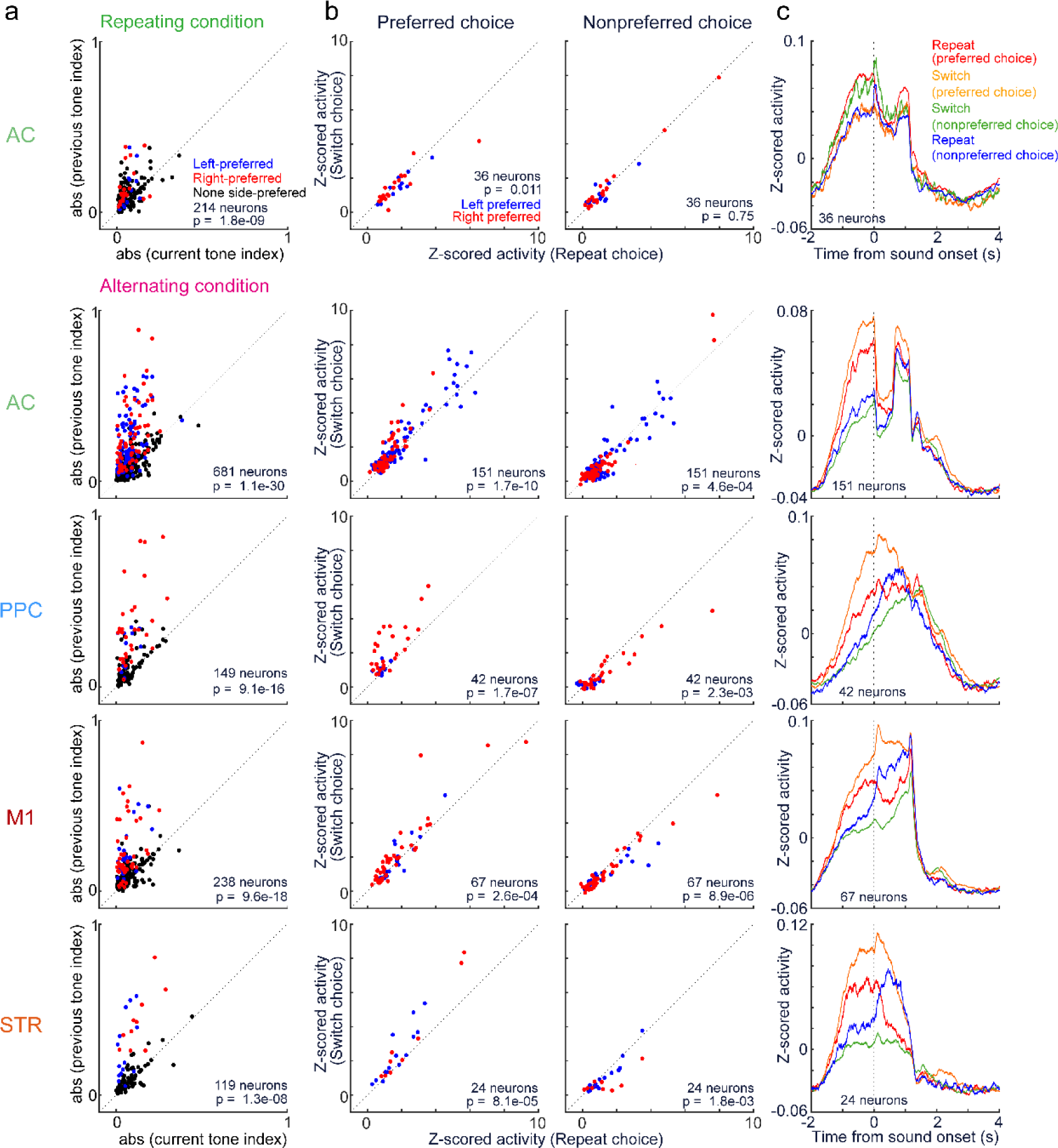
Neural activity in the AC, PPC, M1, and STR before sound presentation. **a.** Scatterplot comparing the absolute tone indices based on the previous and current trials. The previous tone indices were greater than the current tone indices, suggesting that the neurons represented previous choices (blue and red dots: left- and right-side preferred neurons, p < 0.01 in the Mann‒Whitney U test; black dots: non-side-preferred neurons; p value in the Wilcoxon signed rank test). **b.** Comparison of neural activity before sound between the repeated and switched choices (Wilcoxon signed rank test). **c.** Average activity traces of the neurons in **b**. We detected a small number of left- and right-side preferred neurons in the PPC (4 neurons) and HPC (5 neurons) in the repeating condition and in the HPC (5 neurons) in the alternating condition. We did not use the data from these brain regions for analyses.

**Supplementary Fig. 5.**
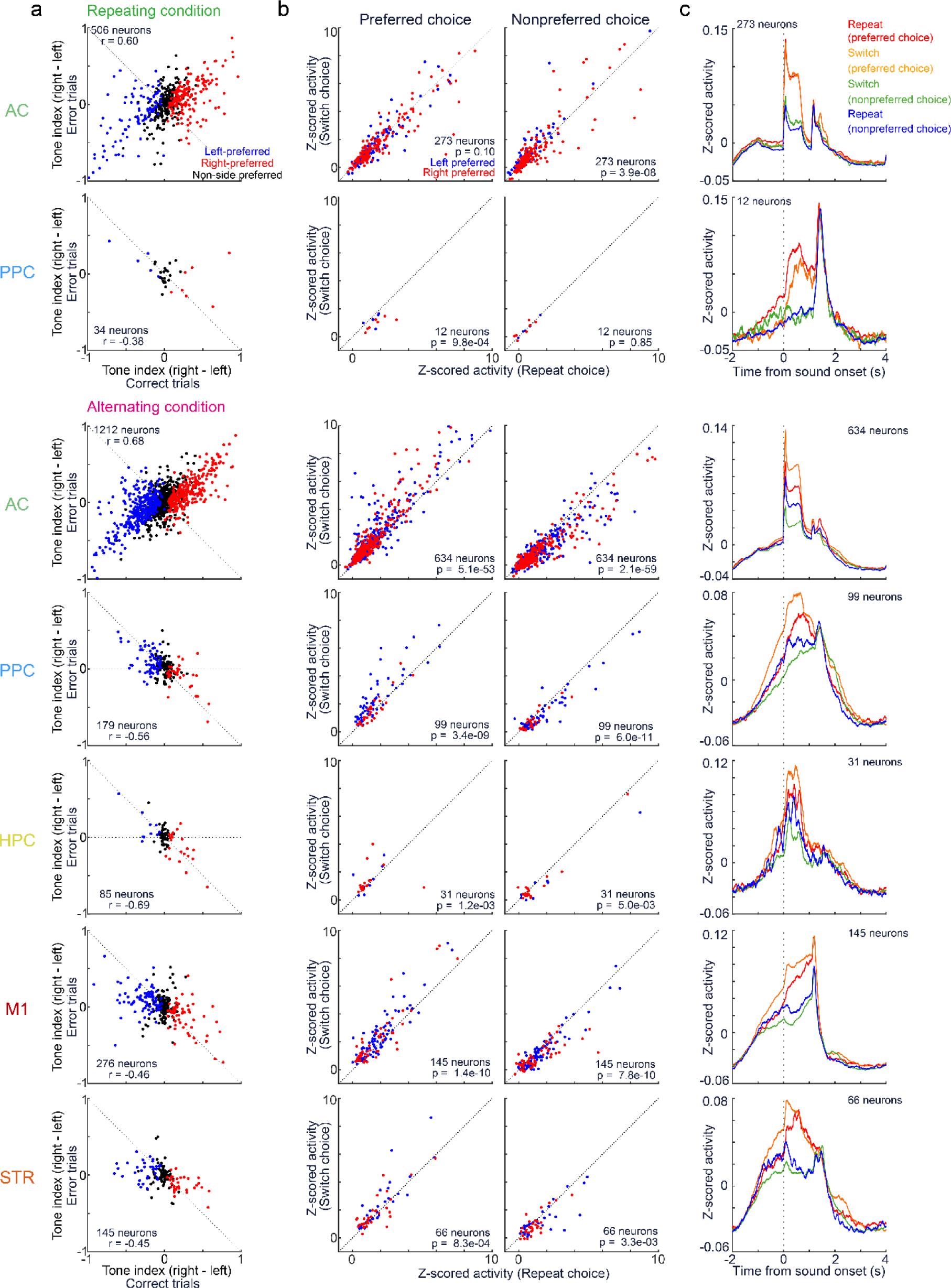
Neural activity in the AC, PPC, HPC, M1 and STR during sound presentation. **a.** Tone index. The neurons in the AC exhibited sound representations, while those in the PPC, HPC, M1, and STR exhibited choice representations. **b.** Comparison of average activity during sound presentation between the repeated and switched choices (Wilcoxon signed rank test). **c.** Average traces of neurons in **b**. In the repeating condition, we did not analyze the neurons in the HPC, as we detected only 2 left- or right-side preferred neurons.

**Supplementary Fig. 6.**
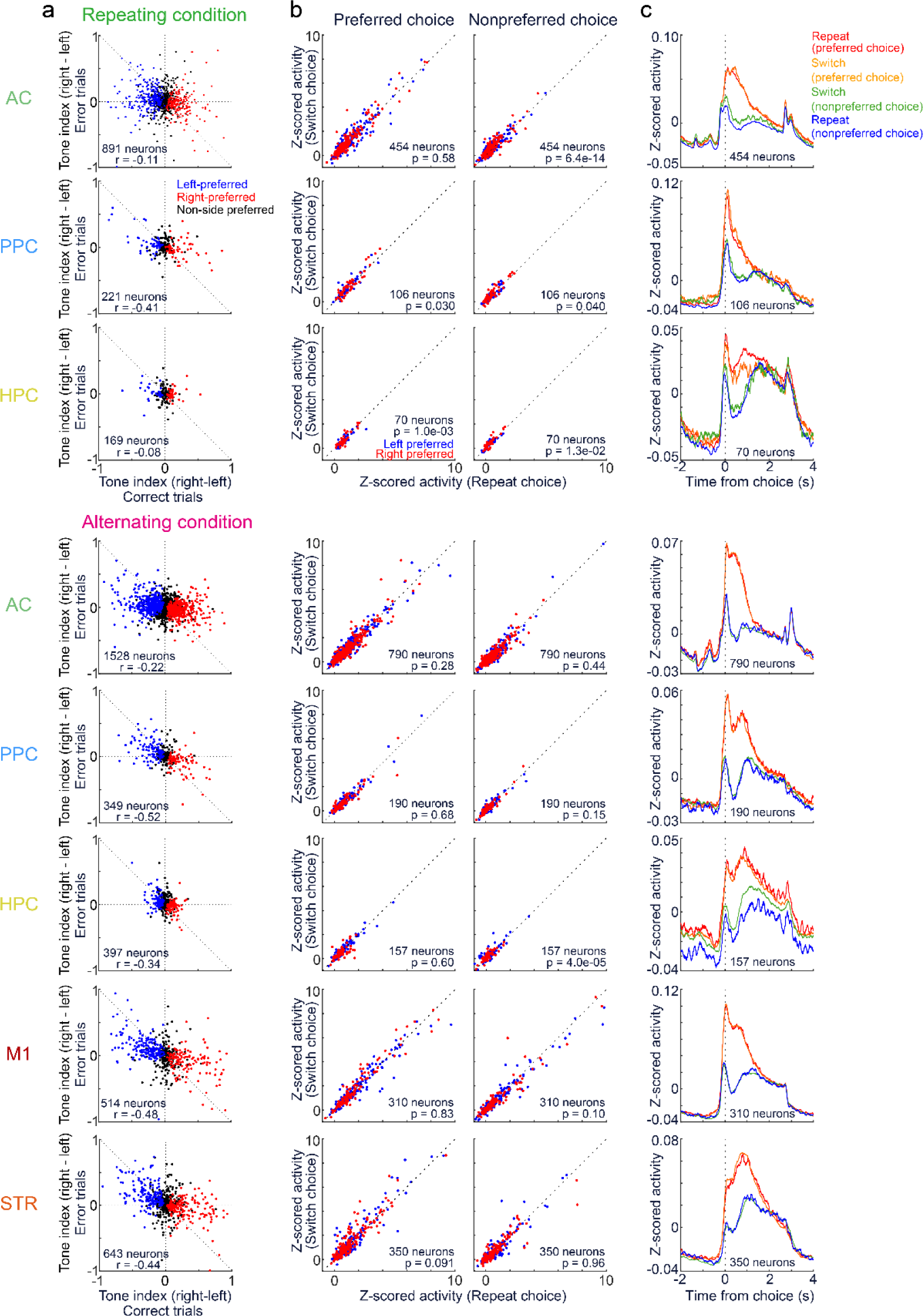
Neural activity in the AC, PPC, HPC, M1, and STR during the outcome in correct trials. **a.** Tone index. The data presentations are the same as those in **Supplementary Fig. 5** but during outcomes and correct choices. The neurons in all the recorded regions represented choices. **b.** Comparison of average activity during 0–1.0 s in correct trials between the repeated and switched choices (Wilcoxon signed rank test). **c.** Average traces of neurons in **b**.

**Supplementary Fig. 7.**
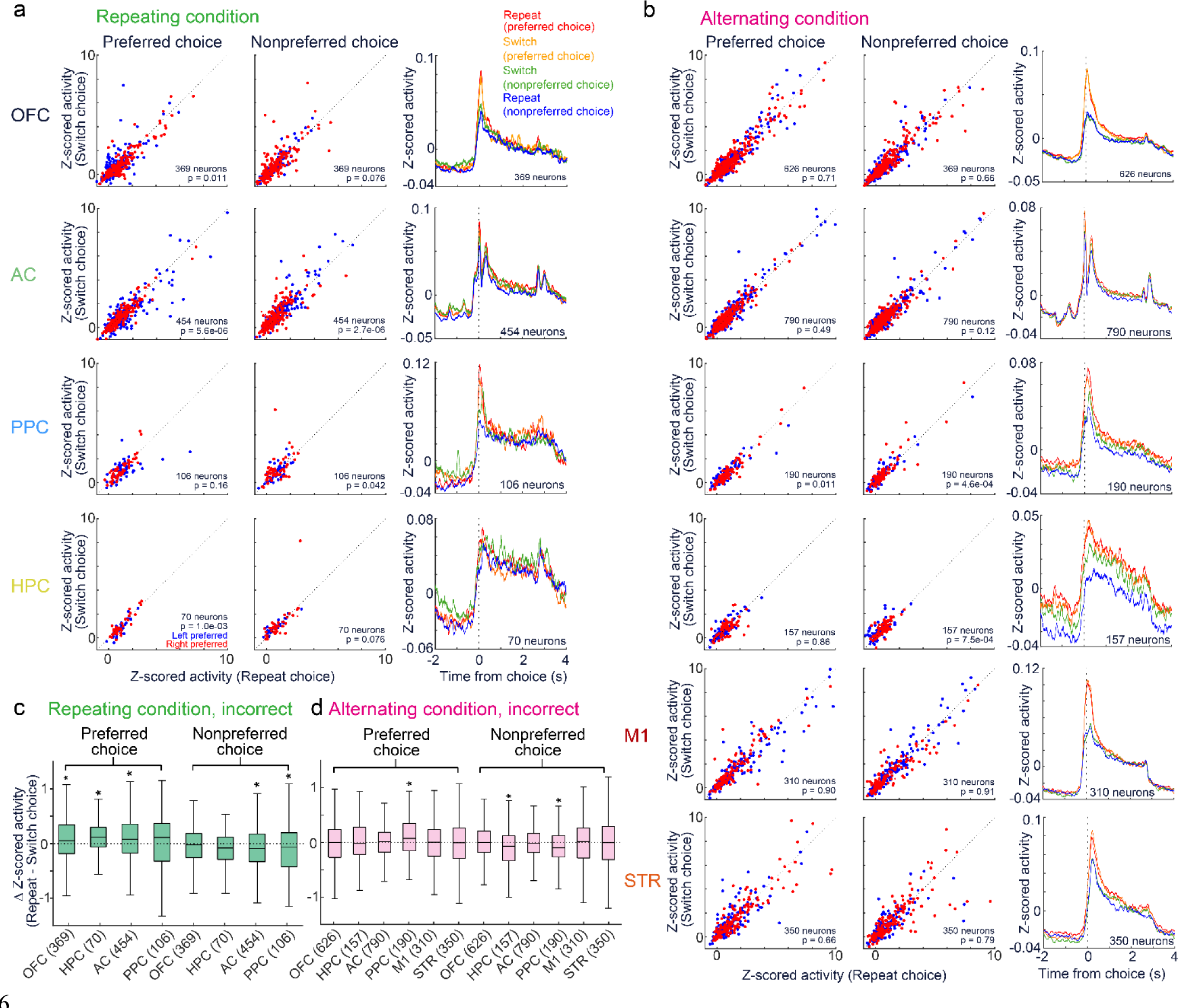
Neural activity in the OFC, AC, PPC, HPC, M1, and STR during the outcome of incorrect trials. **a.** Activity in the repeating condition. The data plots are the same as those in **Supplementary Fig. 6** but for incorrect trials. Left and middle: Comparison of average activity during 0–1.0 s in incorrect trials between the repeated and switched choices (Wilcoxon signed rank test). Right: Average traces of neurons. **b.** Same as **a** but in the alternating condition. **c, d**. The difference in average activity between 0 and 1.0 s from the outcome between the repeated and switched choices in incorrect trials. * p < 0.05 according to the Wilcoxon signed rank test.

